# Infection outcomes depend on within-host spatial dynamics

**DOI:** 10.64898/2025.12.19.695384

**Authors:** Malavika Menon, Daniel Essilfe, Mathias Franz, Sophie A.O Armitage

**Author notes:** Joint last authors (Mathias Franz, Sophie A.O Armitage).

## Abstract

Host-pathogen interactions can result in stochastic variation in infection outcomes, where the host survives the infection with a persistent bacterial load, clears the infection, or succumbs to death with a high pathogen load. This inter-individual variation is thought to be driven by the speed of immune activation, pathogen virulence and the ability of the pathogen to survive inside the host, but not much is known about whether inter-individual variation in infection outcome could stem from other factors, such as within-host spatial dynamics of the pathogen during infection. Hence, in this study, we used the pathogenic bacterium *Providencia burhodogranariea* to investigate the effect of body part and site of injection on infection outcomes in *Drosophila melanogaster*. Although *P. burhodogranariea* disseminated throughout the body as would be expected for organisms with an open circulatory system, the bacteria localized at lower levels in the abdomen compared to the head and thorax, and they were cleared more quickly in the abdomen. Furthermore, based on our empirical results in combination with a theoretical model, we suggest that terminal infections could be a consequence of damage to the thorax. Our results highlight the role that factors such as injection site and body part can influence the spatial dynamics of the pathogen, which can, in turn, drive the infection outcomes.

## 1. Introduction

An infection can result in three major outcomes: 1) a terminal infection leading to rapid death of the host, 2) clearance of an infection, and 3) an infection that persists inside the host (Acuña-Hidalgo et al., 2022; Van Leeuwen et al., 2019; Duneau et al., 2017; Figure 1A). The variability of infection outcomes is observed across various taxa, such as COVID-19 infection in humans, where some individuals showed no or mild symptoms, while others died due to severe symptoms (Pereira et al., 2021), and *Mycoplasma haemomuris* and *Bartonella krasnovi* infection in rodents (Rodríguez-Pastor et al., 2025), and *Plasmodium berghei* infections in *Anopheles stephensi* mosquitoes (Pirahmadi et al., 2025). Such variation in infection outcomes could have ecological and evolutionary implications, such as for pathogen transmission, prevalence, and virulence, i.e., the damage caused by an infection (Acuña-Hidalgo et al., 2022). The variation in infection outcomes is shaped by the interplay of host and pathogen interaction, which includes host genotype (Howick & Lazzaro, 2014), host ecology (Hanson et al., 2023), host immune investment (Lazzaro & Tate, 2022), pathogen virulence (Duneau et al., 2017), and pathogen growth factors (Shaka et al., 2022).

**Figure 1.**
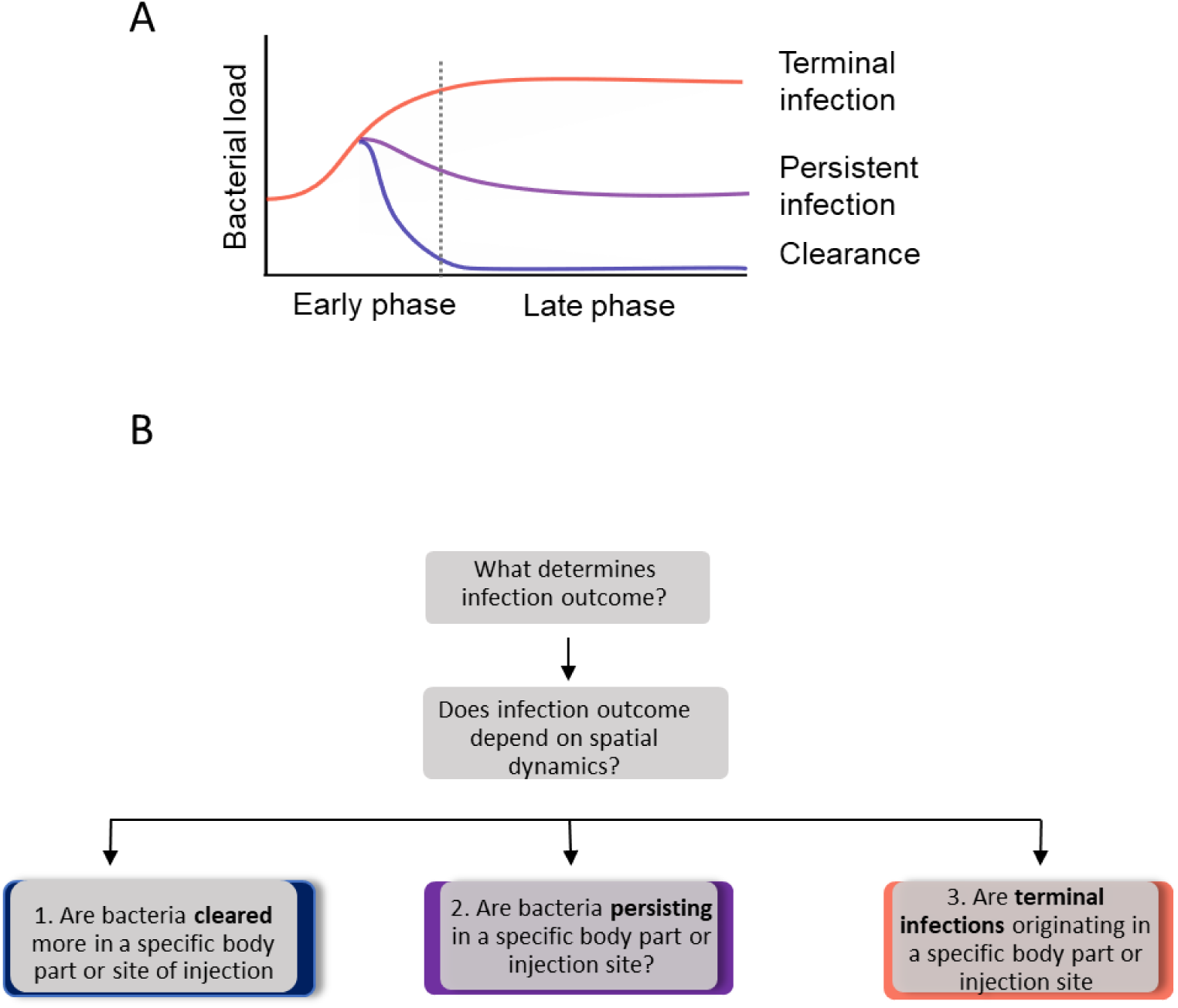
**Possible infection outcomes and the hierarchy of hypotheses for infection outcomes**. A) The three possible infection outcomes visualised in terms of changes in bacterial load over time. Bifurcation in the early phase of infection can lead to high loads and terminal infections, i.e., host death, or it can lead to a persistent infection or clearance of the pathogen (Duneau et al., 2017; Acuña-Hidalgo et al., 2022). B) Illustration for the overall hypothesis for infection outcomes addressed in this paper. Each of the three branches represents one of the infection outcomes: clearance, persistent infection, and terminal infection.

The emergence of infection outcomes could be better understood via an increased understanding of within-host dynamics. Within-host dynamics can be difficult to study in vertebrate models due to ethical concerns; hence, invertebrate models such as *D. melanogaster*, can be used to understand disease progression (Duneau & Ferdy, 2022). A particularly interesting dichotomy in infection outcomes has been documented in experiments in which *Drosophila melanogaster* were experimentally infected with *Providencia rettgeri* and *P. burhogranariea* (Duneau et al., 2017). Despite identical inoculation doses, genetically similar flies followed two opposing bacterial load trajectories and corresponding infection outcomes: either flies exhibited terminal infections with high bacterial loads, which resulted in rapid host death, or flies exhibited persistent infections with considerably lower bacterial loads and without any apparent increase in host death (Duneau et al., 2017; Figure 1A). Duneau et al. (2017) also demonstrated that bifurcation to a terminal or a persistent infection is influenced by variation in the rate of bacterial growth inside the host and the time taken for the immune system to control the infection. Acuña-Hidalgo et al. (2022) subsequently showed that bacterial infection in *D. melanogaster* can also lead to clearance of the infection, meaning that together with terminal- and persistent infections, bacterial loads have the possibility to trifurcate (Figure 1A). However, what causes this intriguing variation in outcomes is currently not well understood, and here we hypothesized that investigating the within-host spatial dynamics of bacterial infections will give insights into this issue.

The variability in infection outcomes was studied using various theoretical models. There seems to be a general agreement that the speed of the host immune reaction after bacterial inoculation is a critical determinant of the resulting infection outcome (Duneau et al., 2017; Ellner et al., 2021; Duneau et al., 2025), a sufficiently fast immune reaction is needed to prevent terminal infection (Duneau et al., 2017; Ellner et al., 2021) or to lead to pathogen clearance (Duneau et al., 2025). However, there are also several fundamental differences among these models, for example, in the question of what enables bacterial persistence. Ellner and colleagues (2021) proposed that bacteria persist in a protected state, which could be a specific tissue in which bacteria are shielded from the host immune response. In contrast, Duneau and colleagues (2025) proposed that persistent infections emerge without the need for a protected state. Accordingly, whether and where bacteria might persist in a protective tissue remains to be assessed empirically.

Several previous studies on *D. melanogaster* and other insects have already provided some fundamental insights into within-host pathogen spatial dynamics. For example, Chambers et al. (2014) demonstrated that the site of infection can strongly influence infection outcome in *D. melanogaster*. They found that thorax injections of *Enterococcus faecalis* and *P. rettgeri* resulted in high bacterial proliferation and host mortality, whereas abdomen injections resulted in lower bacterial loads and minimal mortality (Chambers et al., 2014). Furthermore, wounding and bacterial inoculation in the thorax were found to exacerbate the conditions compared to the abdomen (Chambers et al., 2014).

The infection outcomes can be further determined by site-specific tropism, which refers to the ability of a pathogen to preferentially replicate in a particular host body part or region. During infection, some pathogens, such as *Ehrlichia chaffeensis* and *Mycobacterium marinum,* localize in specific tissues that evade host immunity, such as the hemocytes of *D*.

*melanogaster* (Drolia et al., 2013; Dionne et al., 2003). In addition, Roberts and Longdon (2023) reported that after oral infection with *Drosophila* C Virus, seven *Drosophilidae* species exhibited varying viral loads across different tissues. Most studies have focused on the dissemination of pathogens after oral infection, but in nature, wounding can be frequent (Subasi et al., 2024), and wounds can be entry points for pathogens (Otti et al., 2017). It was found that septic infection with *Pseudomonas aeruginosa* showed varying bacterial load in different segments of the fly, depending on the metabolic state of the bacteria (Hastings et al., 2023). The study demonstrated that the thorax had a higher proportion of regular metabolically active cells, while the abdomen had a higher proportion of persisters (bacteria which transition to a metabolically inactive state under stress) during the later phase of infection (Hastings et al., 2023). These examples highlight the potential importance of considering infections from a within-host spatial perspective and the possibility that some tissues or regions of the body can promote bacterial persistence.

In this study, we first asked whether infection outcome depends on within-host spatial dynamics. We sub-set this overarching question into three separate questions, one for each of the three infection outcomes, i.e., terminal infection, persistent infection, or clearance (Figure 1B). For each question, we also tested whether the site of infection and body part affect the outcome. We asked if 1) the infection is cleared in a specific order based on body part or site of injection 2) the bacteria persist in a specific body part or site of injection and, 3) the terminal infections emerge at the site of injection or specific body part (Figure 1B), and further, we addressed if the terminal infections occurred due to wounding in a specific site.

For these two experiments, we used *D. melanogaster* as a host and *P. burhodogranariea* as the pathogen, a combination which is known to result in these trifurcating infection outcomes (Galac & Lazzaro, 2011; Duneau et al., 2017; Acuña-Hidalgo et al., 2022). Further, using a mathematical model based on Ellner et al. (2021), we explored the potential cause of our observation that bifurcating infection dynamics almost exclusively arise after injections in the thorax.

## 2. Materials & Methods

### 2.1 Experimental organisms

We used the gram-negative bacterium *P. burhodogranariea* strain B isolated from the hemolymph of wild-collected *D. melanogaster,* which is an opportunistic pathogen for *D. melanogaster* (Juneja & Lazzaro, 2009) gifted by Brian P Lazzaro (DSMZ; type strain: DSM-199968).

### 2.2 Fly rearing

The *D. melanogaster* fly population was established from 160 Wolbachia-infected fertilized females, which were collected from Portugal (Martins et al., 2013) and gifted to us by Elio Sucena. The flies were maintained at 25°C on a 12-12 hr light and dark cycle at 70% relative humidity in SYA medium (yeast agar medium), which contains 970 ml water, 100 g brewer’s yeast, 50 g sugar, 15 g agar, 30 ml 10% Nipagin, and 3 ml propionic acid (Bass et al., 2007). Flies were kept in population cages containing approximately 5000 flies with non-overlapping generations. For experimental setups, all flies underwent two rounds of density control (Acuña-Hidalgo et al., 2021). Each round of density control used grape juice agar plates (50 g agar, 600 ml red grape juice, 42.5 ml Nipagin [10% w/v], and 1.1 L water) coated with a semi-liquid paste of baker’s yeast mixed with water. The grape juice agar plate was then kept inside the cage overnight for the flies to lay eggs. The following day plates were taken out from the cage and kept inside the incubator. Larvae in the first instar were picked from the plates and placed into fresh food vials. The larval densities were maintained at 100/vial. For the second generation, the step was again repeated with flies that had emerged and were five days old. The flies, which had emerged, were used to set the egg chamber where a grape juice agar plate was smeared with yeast as mentioned above. The flies laid eggs on the plate, which were then used to pick larvae. This way, the flies were density-controlled during development.

### 2.3 Experimental design for experiment 1

This experiment examined how the spatial distribution of the pathogen is related to each infection outcome: clearance, persistent infection, and terminal infection (Figure 1A). This experiment included two treatment groups, with injection doses of ∼92 CFU and ∼9200 CFU per fly, and an injection control group, where flies were only injected with Ringer’s solution. The flies were assayed 1, 7, or 14 days after injection. The timepoint of day 1 was used to assay infection, as the individuals die quickly compared to days 7 and 14, marking the terminal phase of infection. The bacteria were injected in either the thorax between pteuropleura and mesopleura in the 2^nd^ segment, or between the junction of the dorsal and ventral abdomen in the 3^rd^ segment using fine pulled glass capillaries. The experiment was repeated three times, where 228 flies were injected in each experimental replicate (i.e., eight flies x three body parts x two bacterial dose x three time points x two sites of injection, and for Ringer’s, two flies x three body parts x three time points were homogenized). Injections were carried out in a randomized block design. Flies were dissected and plated for bacterial load as described below.

### 2.4 Experimental design for experiment 2

This experiment aimed to disentangle two co-occurring effects of an injection: 1) the infection starting in a specific body part 2) the infection starting in a specific site due to wounding.

Chambers et al. (2014) observed that injection in the thorax induced acute mortality independent of inoculation. Hence, here we tried to examine if we find a similar pattern during terminal infection. Flies used in the current experiment were maintained and reared as mentioned in the above section. This experiment included two doses, ∼9200 CFU and ∼92 CFU, injected into flies. Here, we used pricking to create wounds without bacterial inoculation. Individual flies were either pricked in the abdomen or thorax or not pricked. Five minutes after pricking, the flies were injected into the thorax or abdomen. This resulted in four treatment groups: 1) thorax injection without abdomen pricking, 2) thorax injection with abdomen pricking, 3) abdomen injection without thorax pricking, and 4) abdomen injection with thorax pricking. One day after injection, the flies were dissected into head, thorax, and abdomen, and plated for bacterial load as described below. The experiment was repeated three times for each dose, where 84 flies were injected for each replicate (i.e., eight flies x three body parts x four injection/pricking combinations, and for Ringer’s, two flies x three body parts) were homogenized. All injections and plating were carried out in a randomized block design.

### 2.5 Bacterial culturing and dose preparation

We used *P. burhodogranariea* strain B stored at 34.4% final concentration glycerol stocks in the -70 °C freezer. We plated the bacteria in Lysogeny agar (LA) plates from glycerol stock aliquots and incubated at 30 °C for 16 hours. Five colony-forming units were picked from the plates and inoculated in 100 mL sterile LB media and incubated overnight at 30 °C at 200 rpm. For dose preparation, we used the protocol from Kutzer and Armitage (2019). The overnight bacterial culture was washed thrice in 45 ml *Drosophila* Ringer’s solution (182 mmol/L KCl; 461 mol/L NaCl; 3 mmol/L CaCl_2_; 10 mmol/L Tris.HCl). The supernatant was removed by centrifugation at 2880 rcf for 10 minutes. Following this, the overnight bacterial suspensions prepared in two flasks are combined. Following this, a 10-fold serial dilution was done three times, and then the optical density of the bacterial solution was measured using an Ultraspec 10 classic (Amersham) at 600 nm for each dilution. The concentration of the bacterial solution was adjusted based on the required infection dose, based on preliminary experiments. OD within a specific range, i.e., 0.1- 0.7, was serially diluted and plated to confirm CFU estimates. Additionally, to confirm the injection dose post-hoc, the specific concentration estimated by OD was serially diluted and plated three times to estimate the CFU. To inject a dose of 92 CFU and 9200 CFU, we adjusted the concentration to 5×10^6^ and 5×10^8^ bacterial cells/mL, respectively. To confirm the concentration, the injected doses were plated on LB medium after serial dilutions from 1-10^6^ as four droplets of 5 µl, and CFUs were counted after 16 hours of incubation at 30 °C.

### 2.6 Fly injections and bacterial load

Injections were carried out in four to five-day-old female flies, which were sexed one day before injection. The flies were anesthetized using CO_2_ for a maximum of five minutes. A total volume of 18.4 nL of the bacterial solution was injected into the thorax of the flies using 0.5 mm, Drummond capillaries, which were pulled to a fine tip using a Narishige PC-10 puller, and ground to a fine point using a Narishige EG-402 grinder. The capillaries were prepared and filled with mineral oil. Injections were carried out using Nanoinject II^TM^ (Drummond). A volume of 18.4 nL accounts for ∼9200 cells from a 5×10^8^ concentration of the bacterial solution and ∼92 cells in a 5×10^6^ concentration of the bacterial solution. Flies were injected at the specific time points to be assessed and kept in groups of six per vial. Following the injection at specific time points, each fly was dissected using fine dissecting scissors. The anesthetized flies were placed on a glass slide under a stereo microscope. The dissection was made in the occipit and the prothorax (first thoracic segment) to separate head and thorax, followed by dissection in the postnotum tergite junction between metathorax and first abdominal segment A1 to separate the thorax and abdomen, which additionally resulted in the dissection of all the adjoining legs along with the thorax. The dissection was done carefully so that the hemolymph did not ooze out. The flies were dissected in the same order every time, and each dissected body part was separately homogenized in 80 µl LB with two beads in a 2 ml microcentrifuge tube kept on ice. The homogenization was done using a Retsch Mill (MM300) at a frequency of 20 Hz for 45 seconds. After homogenization, the fly homogenate was centrifuged at 420 rcf at 4 °C for 1 minute. The fly homogenate was resuspended with a pipette, and 90 µL was transferred into 96-well plates, following which we performed a serial dilution in LB media until 10^6^ and then plated four droplets of 5 µl per fly. The plates were incubated at 30 °C for 16 hours, and CFUs were counted the following day for the dilution where 10-60 CFUs were visible. We did not observe any bacterial growth with Ringer’s injected and naive flies, as *D. melanogaster* microbiota does not grow easily under the above culturing conditions (Acuña-Hidalgo et al., 2022). The lowest detection limit for the bacteria was 1 CFU per body part, which when back-calculated, is equivalent to 5 bacterial cells.

### 2.7 Body size measurement

The volumes of the head, thorax, and abdomen differ from each other; therefore, we used average body part measurements to be able to correct the measured bacterial load for the size of each body part. Using density-controlled flies aged four-days post adult eclosion, we measured the area of the head, thorax, and abdomen for 15 female flies. The flies were placed in lateral orientation. There were two technical replicates for each measurement.

Pictures were taken using 20X magnification with the light microscope (Olympus SZX 16). The area of each body part was measured by using ImageJ by drawing around the body part with the free-hand tool. We found that the head was on average 2.7 times smaller than the abdomen, and the thorax was on average 1.38 times smaller than the abdomen. Accordingly, we multiplied our raw bacterial load measures for head and thorax by the respective factor to obtain body size-corrected load estimates.

### 2.8 Statistical Analysis

The statistical analyses were performed in the statistical software R (R version 4.1.2) (RStudio Team, 2022) using packages “dplyr” (Wickham et al., 2019), “glmmTMB” (Brooks et al., 2017), “car” (Fox & Weisberg, 2019), and “emmeans” (Lenth, 2022). Visualization of the plots was done using “ggplot2” (Wilkinson, 2011). We analysed the bacterial load measurements both with and without the correction for body part size. The results using the correction are presented in the main text, and the results with the raw data are presented in the supplementary information (Figure S1, Table S1, Figure S2, Figure S3).

#### 2.8.1 Experiment 1

##### 2.8.1.1 Order of clearance

We performed two different analyses to investigate whether there is an order in which different body parts tend to be cleared. Here, we subset the data based on clearance (the bacterial load is cleared or is below the detection limit) for all the body parts and across all timepoints (1, 7, and 14 days). In the first analysis, we subset the data based on clearance in the site of injection with respect to the body part. We performed an overall Fisher’s test. In the second analysis, we aimed to assess whether there is a body part that tends to be cleared first. For this purpose, we selected flies that had apparently cleared the bacteria in exactly one body part, assuming that this body part represents the part that was cleared first in these flies. In the second analysis, we aimed to assess whether there is a body part that tends to be cleared last. For this purpose, we selected flies that had apparently cleared the bacteria in two body parts, assuming that the body part that was not cleared yet represents the body part that cleared last in these flies. For both analyses, we first performed an overall and then pairwise Chi-square tests to test whether there was an effect of body part (head, thorax, or abdomen).

##### 2.8.1.2 Spatial variation of bacterial load during the persistent phase of infection

In this analysis, we used a generalized linear mixed model to estimate the bacterial load at 7-and 14-day post-infection, i.e., during the persistent phase of the infection. As a response variable, we used double-log transformed bacterial load. The fixed effects included body part, site of infection, and time point. The random effects included replicate and fly id. In addition, to prevent violation of model assumptions, we assumed that all considered fixed effects can also affect the residual variance (which we implemented using dispformula in the function glmmTMB).

Model 1: log (log (bacterial load)) ∼ body part + site of injection + dose + timepoint + (1|replicate) + (1|fly id)

##### 2.8.1.3 Occurrence of terminal infection

In this analysis, we used data on bacterial load 1-day post-infection to investigate 1) whether the occurrence of terminal infections depends on the site of infection, and 2) whether in thorax injections, terminal infections tend to originate in a specific body part. To address the first question, for each fly, we obtained the total bacterial load by summing the measured loads of each body part. Each fly was classified as having either a “low load”, which most likely would result in clearance or persistent infection, or a “high load”, which would most likely lead to a terminal infection. As a cut-off for this classification, we assumed a total load of 10 times the injection dose to be included in the “high load” group. We then used Fisher’s exact test to assess whether the site of injection is related to the occurrence of high loads. To address the second question, we used only flies that were injected in the thorax and had not cleared the infection. We separately classified each body part as “low load” or “high load” based on whether the measured load in this body part was larger than 10 times the injection dose. We further used Fisher’s exact test to assess a potential relationship between body part and the occurrence of high loads.

#### 2.8.2 Experiment 2

##### 2.8.2.1 Occurrence of terminal infection due to wounding in the thorax

In this analysis, we used only day 1 post-injection to decompose the effect of sterile wounding and bacterial inoculation in the thorax on the occurrence of terminal infection. The data consists of body part bacterial load in four treatments: 1) abdomen pricking-thorax injection, 2) no pricking abdomen injection, 3) no pricking thorax injection, and 4) thorax pricking-abdomen injection. In this analysis, we address whether terminal infection in the thorax occurs due to wounding or pathogen inoculation in the thorax. We used threshold dose*10 to test the difference in high loads and low loads among the treatments and body parts using Fisher’s test.

### 2.9 Theoretical model

We developed a theoretical model to explore the potential cause for our observation that bifurcating infection dynamics almost exclusively arise after injections in the thorax. For this purpose, we modified the model developed by Ellner et al. (2021). The key mechanism for bifurcating infection dynamics in the models of Ellner et al. (2021) includes (1) the production of bacterial proteases that degrade host AMPs, and (2) the sequestration of AMPs during the killing of bacteria. When the bacterial population is sufficiently large, these mechanisms successfully counter the host’s immune defence and lead to uncontrolled bacterial growth. To allow the emergence of persistent infections, in one of their models, Ellner and colleagues included the possibility that bacteria can enter a protected state, where they are protected from the host immune response.

Here, we investigated a variation of this protected state model. In contrast to Ellner et al. (2021), our main interest here was not the emergence of persistent infections. Instead, we hypothesized that a protective tissue that mainly occurs in the thorax, e.g., due to damage of wing muscles, could cause the emergence of terminal infections. Therefore, in our model, we changed several assumptions. Most importantly, we assumed that bacteria in the protective tissue grow at the same rate as bacteria in the hemocoel. Additionally, we assumed that bacteria enter and leave the protective tissue independent of hemocoel AMP concentrations, and bacteria in the protective tissue do not release proteases into the hemocoel.

Our model is a direct extension of the model described in equation 2.2 in Ellner et al. (2021) and contains three state variables: bacteria *B* in the hemocoel, bacteria *P* in the protected tissue, and scaled AMPs *A* in the hemocoel:

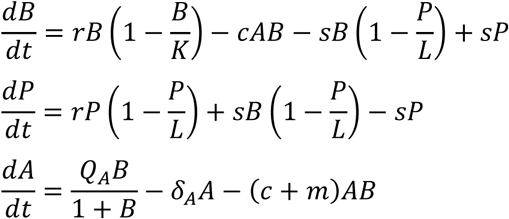

Bacteria grow at a rate *r* towards the carrying capacities *K* in the hemocoel and *L* in the protected tissue. In the hemocoel, bacteria are killed by host AMPs at a rate *c*. Transitions of bacteria between the hemocoel and protected tissue occur at rate *s*. For transitions from the larger hemocoel to the smaller protected tissue the transitions are additionally assumed to follow a logistic function so that transitions to the protected tissue decrease as the bacterial population size in the protected tissue approaches the corresponding carrying capacity *L*. Dynamic of AMPs in the hemocoel follow the assumptions of Ellner and colleagues: AMPs production follows a Hill function with a maximum AMP production rate of *Q_A_*, AMPs are naturally degraded with rate *δ_A_*, and AMPs are degraded (1) by sequestration to killed bacterial with rate *c* and (2) by bacterial proteases with rate *m*.

The main aim of our model analysis was to provide a proof of concept that variation in protected tissue size is a so far unrecognized possibility for the generation of bifurcating infection dynamics. For this purpose, we used the same parameter values as Ellner et al. (2021) in their Figure 3, i.e., *r* = 0.5, *K* = 1000, *c* = 0.1, *Q_A_* = 10, *δ_A_*= 0.02, and *m* = 0.2. Additionally, we assumed *s* = 0.2 and investigated the following values for the carrying capacity of the protected tissue *L*: 2, 5, 10, 20, 50, and 100. We assumed that such differences in *L* can be caused by different amounts of damage to wing muscles in the thorax during the injection of bacteria. In our simulations, we assumed initial conditions of *B_0_* = 1, *P_0_* = 0, *A_0_*= 0, and simulated dynamics for 100 hours.

## 3. Results

### 3.1 Pathogen clearance starts more frequently in the abdomen

We found that 82 individuals out of 376 individuals showed clearance, among which 28 individuals showed complete clearance in all the body parts, 30 cleared in only one body part, and 24 cleared in two body parts. We started by testing the effect of the site of injection with respect to the body part for clearance. We found no significant difference based on the site of injection across different body parts for the first to clear. Further, we tested which body part the pathogen was cleared from first or last. There was a significant overall effect of body part on the frequency of the body part that was first to be cleared (*χ^2^* = 10.400, df = 2, *p* = 0.005; Figure 2A). Pair-wise comparisons between the body parts showed that the abdomen was more likely to be cleared before the head (*χ^2^* = 3.846, df = 1, *p* = 0.049) and the thorax (*χ^2^* = 8.909, df = 1, *p* = 0.003), whereas there was no apparent difference between the head and the thorax (*χ^2^* = 1.333, df = 1, *p* = 0.248). In contrast to the first to be cleared, we found no apparent effect of body part on the frequency of the body part that was last to be cleared (*χ^2^* = 0.750, df = 2, *p* = 0.687; Figure 2B). Additionally, we also found no difference across the site of infection with respect to clearance in the body parts for the last to clear (*p* = 0.615).

**Figure 2.**
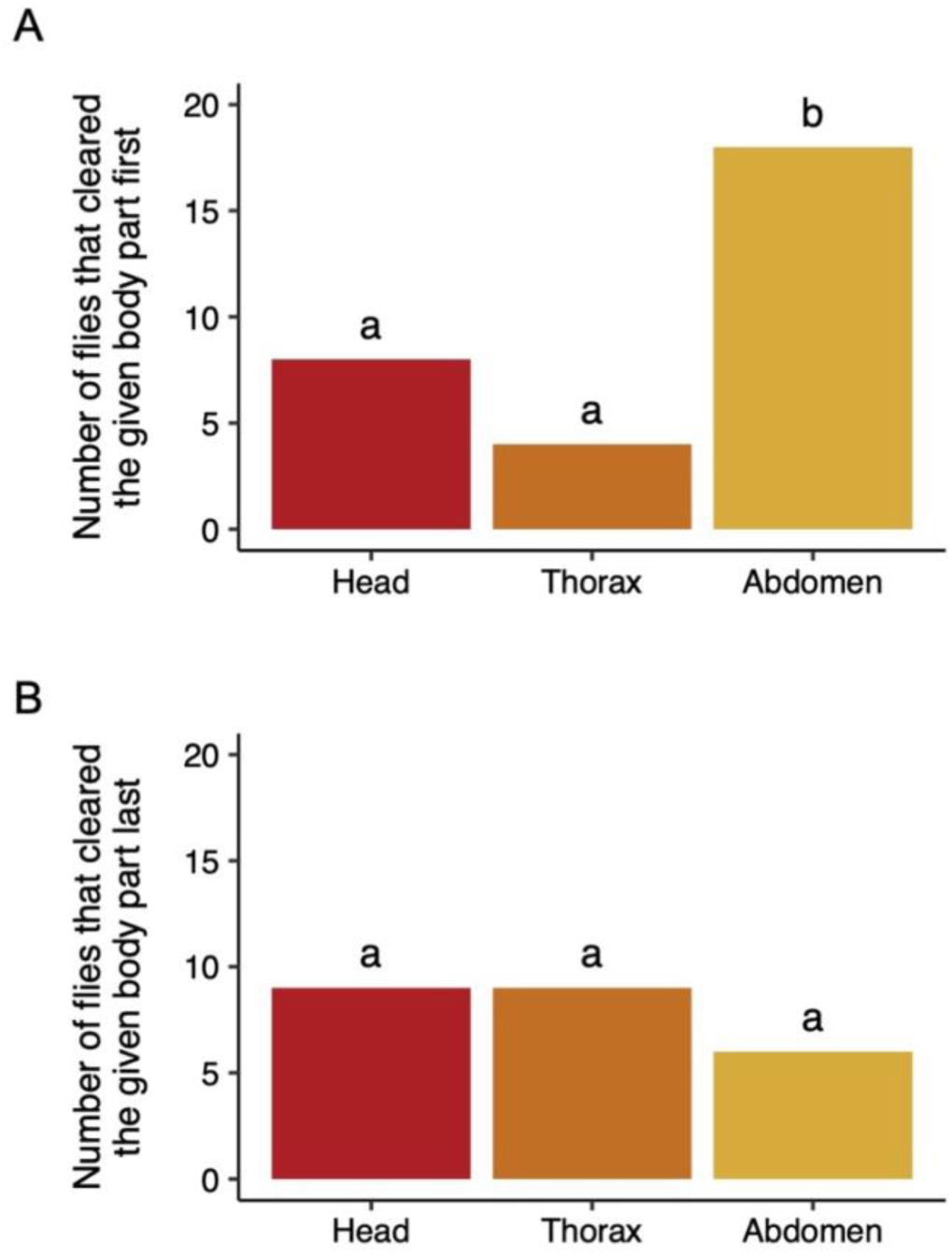
Order of pathogen clearance in the different body parts. A) First body part to be cleared. The data set includes only flies that had cleared exactly one body part (n = 30 flies out of a total of 376 flies). Here, the abdomen was more frequently the first to be cleared compared to the head and thorax. B) Last body part to be cleared. The data set includes only flies that had cleared exactly two body parts (n = 24 flies out of a total of 376 flies). Here, there was no indication of differences among body parts. Data sets in both panels include both injection doses and injection sites, and flies were homogenized 1, 7, and 14 days after injection. Where letters above the body parts are not the same, the frequency of body parts is significantly different from each other.

**Figure 3.**
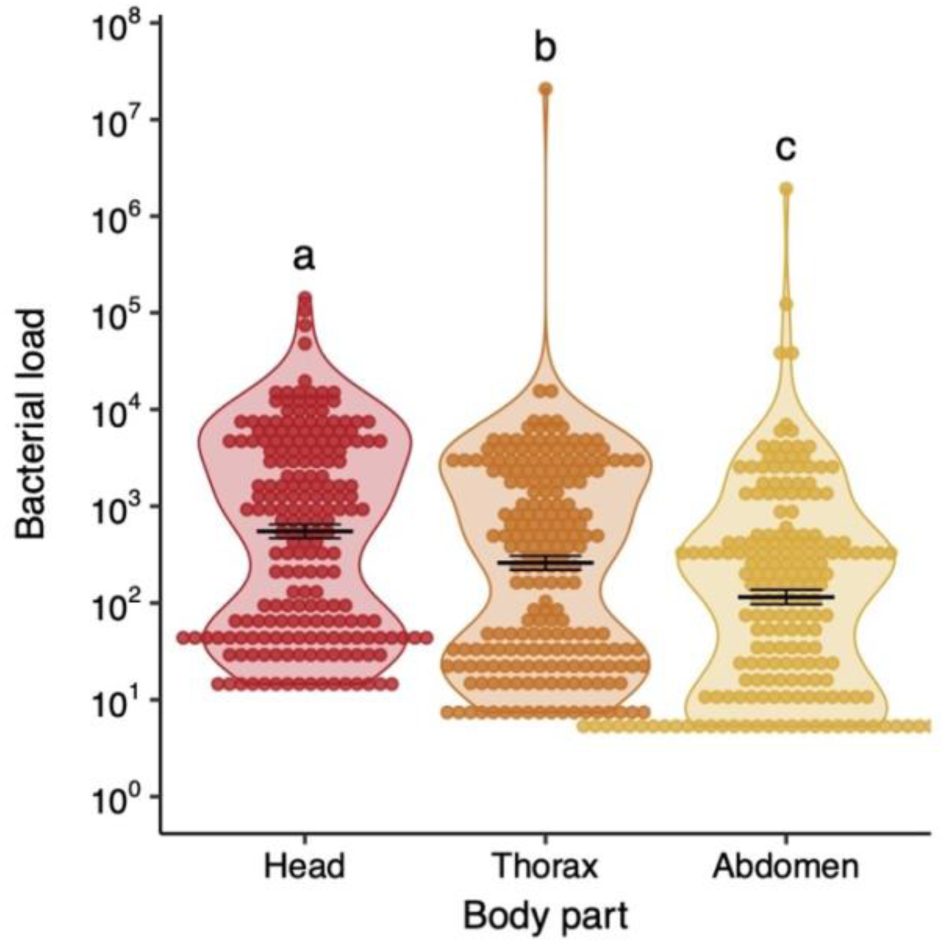
**Bacterial load across body parts during persistent infection**. The figure includes both injection doses and injection sites, and flies homogenised 7- and 14-days after injection, i.e., only the persistent infection phase. Flies that had completely cleared the infection were not included. Each data point is the body part of one fly, and the black lines indicate the arithmetic means. Where letters above the treatment groups are not the same, the treatment groups are significantly different from each other. Sample sizes for each group are: head n = 204, thorax n = 207, abdomen n = 193.

### 3.2 The abdomen sustains lower loads during persistent infection

We assessed all flies with non-zero bacterial loads in different body parts at 7- and 14-day post-injection. Bacterial load was not affected by of site of injection. However, there was a significant effect of body parts, dose, timepoint, and replicate (Figure 3, Table 1). In this study, we focus only on body parts. Post-hoc tests revealed that there is a significant difference between head vs thorax, thorax vs abdomen, and head vs abdomen, where head displayed a higher bacterial load compared to thorax and abdomen, while the abdomen displayed a lower load compared to head and thorax (Table 2). In an additional experiment, we tested the generality of the persistent phase of infection in males and females. We found a similar pattern in bacterial load for both sexes; the head had a higher bacterial load compared to the thorax and abdomen, and the thorax showed a higher bacterial load compared to the abdomen. This suggests that the abdomen sustains a lower bacterial load irrespective of the sexes (Figure S2).

**Table 1.** Results of a linear mixed model with bacterial load on days 7 and 14 as the response variable. Statistically significant P values are in bold.

| *Factor* | *Chisq* | *df* | *P value* |
| --- | --- | --- | --- |
| Body part | 263.359 | 3 | **< 0.001** |
| Site of injection | 2.973 | 1 | 0.084 |
| Dose | 1093.295 | 1 | **< 0.001** |
| Timepoint | 39.024 | 1 | **< 0.001** |
| Replicate | 70.989 | 4 | **< 0.001** |

**Table 2.**
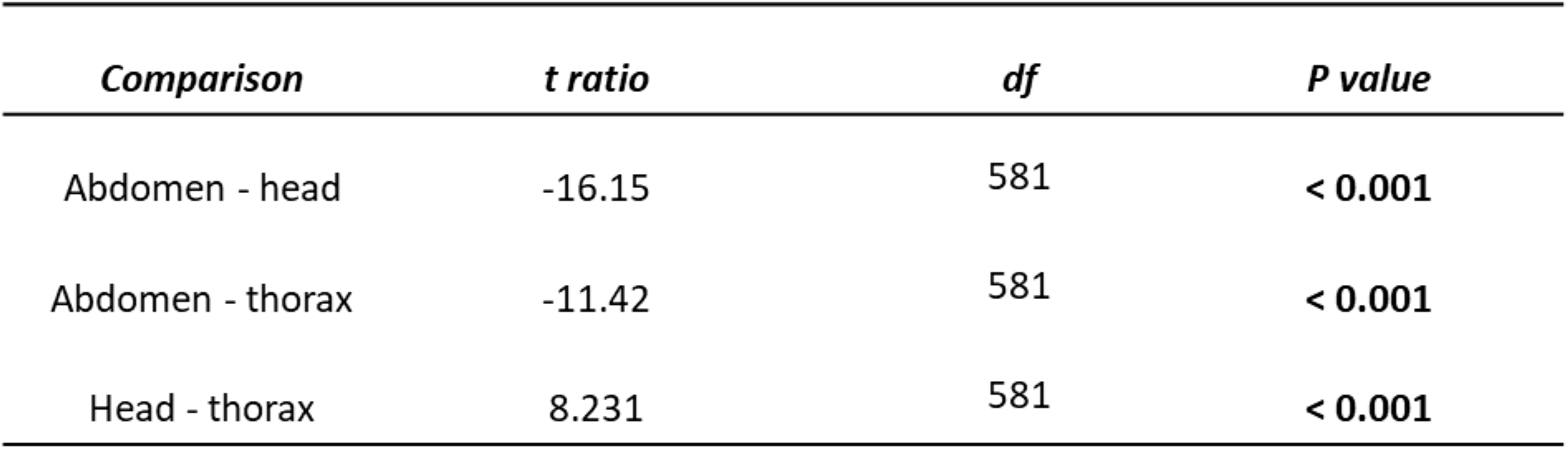
Tukey post-hoc tests for body part comparisons. Statistically significant P values are in bold.

| *Comparison* | *t ratio* | *df* | *P value* |
| --- | --- | --- | --- |
| Abdomen - head | -16.15 | 581 | **< 0.001** |
| Abdomen - thorax | -11.42 | 581 | **< 0.001** |
| Head - thorax | 8.231 | 581 | **< 0.001** |

### 3.3 Damage to the thorax results in the occurrence of terminal infections

In the first experiment examining terminal infections, we tested whether specific body parts or the site of injection influenced the occurrence of terminal infections at 1-day post-injection.

When summing up total bacterial load in whole flies, we found that thorax injections resulted in a larger proportion of individuals with high loads, i.e., terminal infections, compared to abdomen injections (*p* < 0.001; Figure 4A & S3).

**Figure 4.**
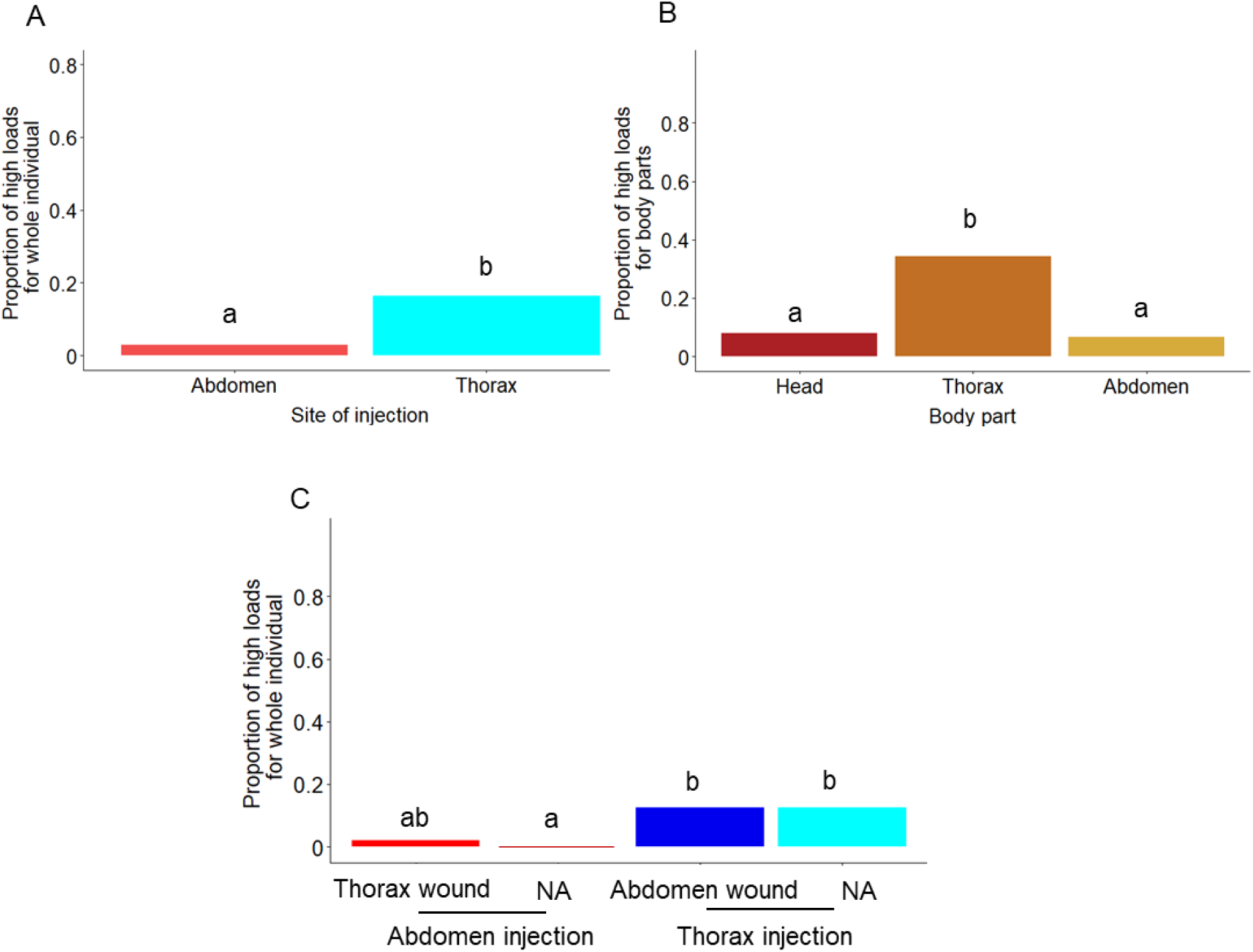
**The occurrence and origin of terminal infections**. Experiment 1: A) The effect of site of injection on the proportion of flies with high total loads, i.e., the bacterial loads were summed up across the head, thorax, and abdomen of each fly. Sample sizes are: abdomen injection, n = 41 out of 123 individuals; thorax injection, n = 75 out of 123 individuals. B) The effect of body part on the proportion of flies with high total loads. Sample sizes are: head, n = 75 ; thorax, n = 76 ; abdomen, n = 74 . Panels A and B include both injection doses and flies homogenised 1-day post-injection, i.e., only the early infection stage. Experiment 2: C) The effect of wounding and site of injection (route of pathogen entry) on the proportion of flies with high total loads. The flies were injected with 92 CFU and 9200 CFU, and they were homogenised one day post-injection, i.e., only the early infection phase. Where letters above the treatment groups are not the same, the treatment groups are significantly different from each other. The total sample size for high loads in all treatments is n = 48 out of 192 individuals.

For thorax injections, we analysed the occurrence of high loads in each body part. In this analysis, we found that the occurrence of high loads differed among body parts (*p* < 0.001; Figure 4B). Specifically, we found a higher proportion of individuals having high loads in the thorax compared to the abdomen (*p* < 0.001) and the head (*p* < 0.001), whereas we found no difference in the head compared to the abdomen (*p* = 0.651).

In the second experiment examining terminal infections, we tested whether the effect of injection site and body part observed in the first experiment (Figure 4A & B) was caused by wounding and whether wounding in the thorax increases the occurrence of terminal infections independently of the site of bacterial injection. We found that the occurrence of high total loads differed significantly across the four treatment groups (*p* = 0.003, Figure 4C). In the absence of wounding, we observed similar results to the first experiment, i.e., high loads were more frequent in thorax injections compared to abdomen injections (*p* = 0.015). For all the other pairwise comparisons, we did not find statistically significant differences (Table S5), except for thorax injection combined with abdomen wounding, which resulted in more frequent high loads compared to abdomen injection without wounding (*p* = 0.015).

Therefore, these results do not support the idea that wounding in the thorax increases the occurrence of terminal infections independently of the site of bacterial injection. However, note that in this experiment, we observed a generally low proportion of high loads, which reduced statistical power

### 3.4 Bifurcating infection dynamics in the thorax

Our simulated results confirm that variation in protected tissue sizes is a potential cause for the generation of bifurcating infection dynamics (Figure 5). Specifically, infections are controlled at low bacterial loads by the host immune response for smaller carrying capacities of the protected tissue *L*, but infections grow out of control for higher bacterial loads with larger carrying capacities. These dynamics emerge because after the initial colonization of the protective tissue, this tissue acts as a source of the population, with more bacteria transitioning from the protective tissue to the hemocoel than the other way around (Figure S4). The bacteria in the protected tissue can grow without the harmful effects of the host’s immune response. Bacterial transitions from the protected tissue to the hemocoel cause an increased growth of the bacterial population in the hemocoel. Importantly, the number of bacteria that transition from the protected tissue to the hemocoel increases with increasing carrying capacity of the protected tissue (Figure S4). If this additional inflow of bacteria to the haemocoel becomes sufficiently strong, then this overwhelms the host immune response and the bacterial population in the hemocoel grows out of control (Figure 5). The bifurcating infection dynamics appear to be robust to other parameters such as transition rate *s* (Figure S5).

**Figure 5.**
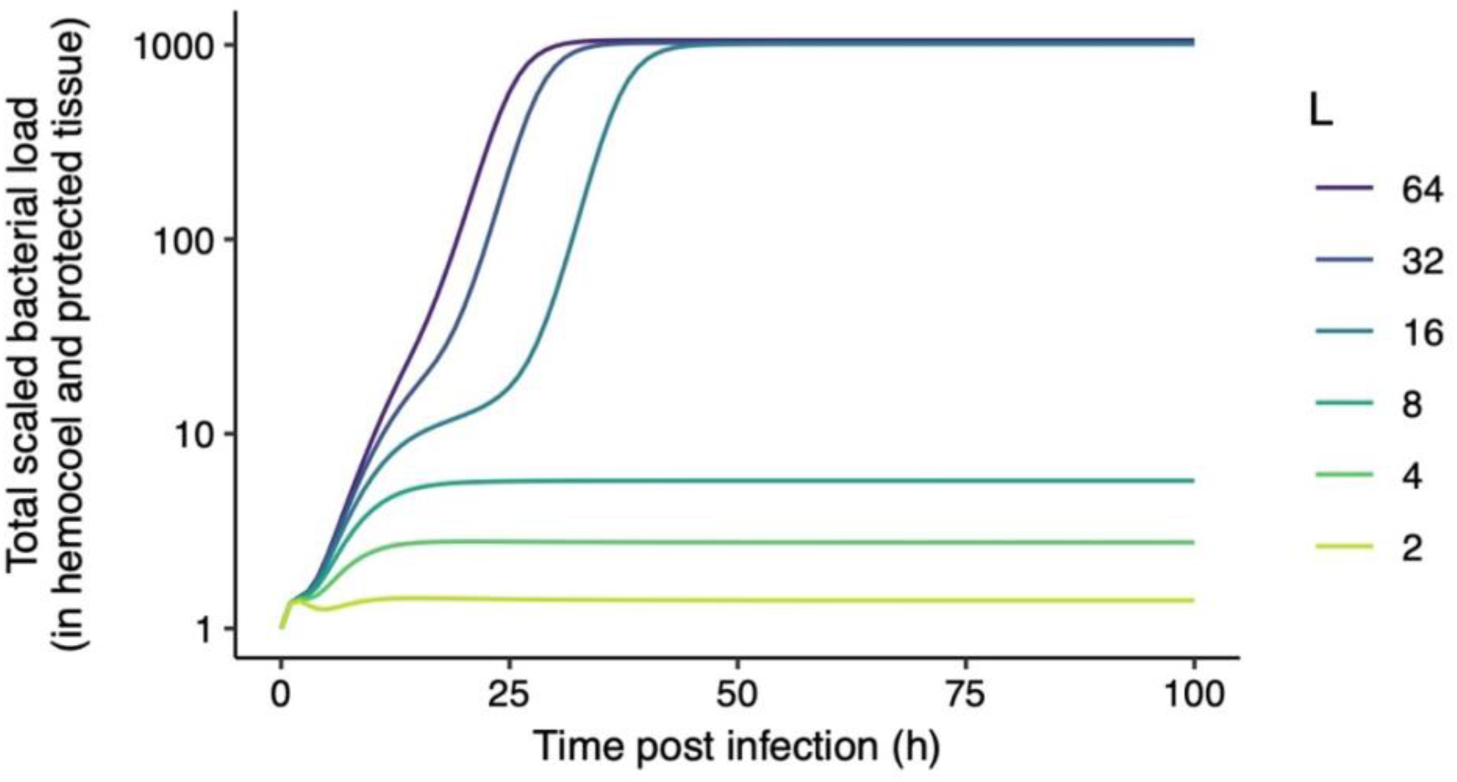
Simulated model dynamics depicting temporal changes in total bacterial population load for different carrying capacities of the protected tissue *L*. The different colours represent the varying values of the carrying capacity of the protective tissue *L*, which reflect different limits of bacterial load up to which bacteria can still grow in the protective tissue. At low values of *L*, bacteria are suppressed by the immune response in the hemocoel and are therefore largely restricted to the protective tissue. Accordingly, *L* closely matches the total bacterial load in the whole fly. In contrast, if *L* is sufficiently high, bacteria can always grow to a similar, very high total load. This switch in the bacterial growth dynamics emerges because higher *L* leads to an increased number of bacteria that transition from the protected tissue to the hemocoel (Figure S4). A sufficiently strong inflow of bacteria into the hemocoel eventually overwhelms the host’s immune response, and the bacterial population in the hemocoel grows out of control.

## 4. Discussion

In this study, we aimed to understand within-host pathogen spatial dynamics and their relationship to infection outcomes. We found that infections were cleared first in the abdomen. Second, during the persistent infection phase, although *P. burhodogranariea* disseminated throughout the host, it had lower loads in the abdomen compared to the head and thorax, which is in line with the results on clearance. Third, we found a higher occurrence of terminal infections in the thorax compared to the head and abdomen. However, when we attempted to disentangle the effects of wounding and pathogen injection at a specific site, we found no conclusive evidence that injury in the thorax was solely responsible for terminal infections, due to difficulties in obtaining flies with terminal infections. Nevertheless, the results of our theoretical model support the idea that wounding can facilitate the emergence of terminal infections if wounds allow bacteria to enter a tissue where they are protected from the host immune response.

### 4.1 Pathogen clearance starts more frequently in the abdomen

The tendency of *P. burhodogranariea* to be cleared first from the abdomen (Figure 2A) before the head and thorax could be due to several reasons, for example, due to differences in immune activity or anatomy between the body parts (Tsou et al., 2000; Schlomann et al., 2024). When a pathogen enters the insect hemolymph, periostial hemocytes localized along the heart of the Drosophila are immediately recruited to the site of infection (Meyer et al., 2023). The periostial immune response most likely occurs in areas where the haemolymph flow is high, regions most likely to encounter pathogens, such as the abdomen (Sigle & Hillyer, 2016; King & Hillyer, 2012; Subasi et al., 2024). The abdomen is also the major site of the fat body, which are associated with the production of antimicrobial peptides responsible for the innate immune response (Meister et al.,1997; Lemaitre & Hoffmann, 2007). The faster clearance in the abdomen compared to other body parts may result from a higher likelihood that AMPs and haemocytes reach the invading pathogen faster in the abdomen compared to the other regions. The faster action of AMPs and the hemocyte recruitment could also be facilitated by the flow of hemolymph current from the posterior to the anterior end (Rotstein & Paululat, 2016; Wasserthal, 2007). The infection-mediated response is likely induced by a combination of circulatory currents and immune components (Yan & Hillyer, 2020). We found no difference in the body part for the last to clear the infection (Figure 2B). This could be due to systemic spread of AMPs and hemocytes through the hemocoel. Once the immune system is activated, it might eventually exert a similar response throughout the body to resist pathogen load, thereby resulting in no specific order for the last to clear.

### 4.2 The abdomen sustains lower loads during persistent infection

A previous study by Robert and longdon (2023) reported that viral infection in different species of *Drosophila* was broadly consistent across its various tissues. Consistent with this study, we found that the pathogen *P. burhodogranariea* was recovered from all body parts; however, the bacterial load differed across body parts. We found a lower bacterial load in the abdomen when compared to the head and thorax. These results were consistent in both corrected and uncorrected bacterial load data (Figure 3, S1A, S1B, Table S1 & S2). The pattern of a lower load in the abdomen compared to the two other body parts was consistent across females and males (Figure S2), suggesting that a general mechanism could be responsible for this finding across the sexes. The lower load in the abdomen complements the faster clearance in the abdomen. Our findings are not consistent with previous reports by Hastings et al. (2023), where they showed that infection of *D. melanogaster* injected in the thorax with regular cells of *P. aeruginosa* resulted in higher loads in the thorax and lower loads in the head during the later phase of infection. The lower load in the head was suggested to be due to a smaller size and minimal circulation of hemolymph in the head compared to the thorax and abdomen. Although it is to be noted that in our study, we only found a higher bacterial load in the head compared to the thorax when corrected for body size. However, in both methods of analysis, we find that the abdomen has the lowest bacterial load compared to the other two body parts. Differences between our findings and those of Hastings et al. (2023) could also be due to the nature of the pathogen because pathogens like *P. aeruginosa* are known to change to a persister state, and the head might be enabling a micro-environment for transitions to the persister state. The pathogen in our study could also be entering a persister state, resulting in lower bacterial detection, however, *P. burhodogranariea* is not well studied for its biofilm and persister ability *in vivo* (Galac & Lazzaro, 2011). The increased prevalence of the pathogen in the head could be due to the microenvironment, which assists in pathogen proliferation or establishes a protective niche that helps to evade the host immunity. Furthermore, *D. melanogaster* possesses a hemolymph-brain barrier that separates hemolymph from neural tissues in the head (Winkler et al., 2025). This blood-brain barrier might prevent pathogen entry into the brain; some bacteria, such as *Streptococcus agalacticae*, *Streptococcus pneumoniae*, *Listeria monocytogenes* etc, could bypass the barrier in *Drosophila* larvae (Benmimoun et al., 2020), although it is unknown if this is the case for *P*. *burhodogranariea*.

Since the site of infection influences the immune response (Nehme et al., 2007; Chen et al., 2024), we expected that the injection site could result in varying localization of the pathogen. However, although we found differences in bacterial load across body parts, we found no difference based on the site of injection. Most studies compared bacterial spread in oral and septic infection (Nehme et al., 2007; Chen et al., 2024). As it is observed notably in mosquito adults that infection induces hemocyte aggregation in the periosteal region, which predominantly encompasses the thorax and abdomen, this can result in a favourable microenvironment for pathogen proliferation (Sigle & Hillyer, 2016). Although in this study we do not find evidence for a favourable microenvironment for the pathogen at the site of injection, it could be that the host immune system is activated irrespective of the site of injection.

### 4.3 Damage to the thorax results in the occurrence of terminal infections

Terminal infections are characterized by a high bacterial load in the host, where ultimately the host succumbs to death. The higher proliferation of bacteria could be a consequence of better nutrient availability, evasion of host immunity within a specific body part, or the pathogen gaining a head start at the site of injection (Dionne et al., 2006; Chambers et al., 2014; McGonigle et al., 2016). We found that terminal infections occur more frequently in the thorax compared to the abdomen (Figure 4A & B). Similar observations were also found by Chambers et al. (2014) where thorax injections with *P. rettgeri* cause increased pathogen proliferation and higher mortality in the *D. melanogaster* (Chambers et al., 2014). The increased susceptibility of flies to thorax injection could be the result of damage to the wing muscles, which are crucial for the flies, resulting in increased investment to repair the damage rather than to resist infection (Apidianakis et al., 2007).

In our second experiment, we aimed to disentangle the effects of wounding and bacterial inoculation from each other. Similar to the first experiment, in the absence of additional wounding, terminal infections were more frequently found after thorax injections compared to abdomen injections. However, when wounding in the abdomen was combined with thorax injection and vice versa, we found no effect of site of injection, as there was no significant difference between wounding in the thorax along with abdomen injection and abdomen injection alone (Figure 4C). This suggests that wounding in the thorax alone does not exacerbate infections. Our results are contrary to the findings by Chambers et al. (2014), who reported a significant difference in wounding in the thorax coupled with abdomen injection compared to abdomen injections alone. This difference might stem from the bacteria, as Chambers et al. (2014) have used *P. rettgeri,* which, although an intermediate pathogen like *P. burhdogranariea,* shows a large variation in bacterial load from 10^3^ to 10^7^ CFU after 24 hours post-infection, while *P. burhodogranariea* maintained stable bacterial loads of 10^3^ to 10^4^ CFU (Galac & Lazzaro, 2011). The large variation in bacterial load might reflect the higher proportion of individuals with higher load *in P. rettgeri* in Chambers et al., (2014). Interestingly, it has been found that terminal infections can also occur after abdomen injection (Duneau et al., 2017). We cannot conclude that the wounding due to a thorax injection is solely responsible for terminal infection. It could be that wounding *per se* and not specifically wounding of the wing muscles in the thorax, leads to terminal infections. Another alternative explanation would be that wounding in the thorax alone is not sufficient to cause terminal infection and needs to be coupled with bacterial inoculation.

Although our empirical evidence suggests there is no role of a protective tissue, our theoretical model, in combination with the empirical results, suggests a potential role of protective tissue in the emergence of terminal infection. Our model suggests that injury in the thorax could result in bacteria entering protected tissues; the variation in protected tissue reflects the different carrying capacities and transition rate (Figure 5 & S5). Our model was able to replicate the bifurcating infection dynamics with both these parameters.

As the transition increases from the bacteria in the protected state to the extracellular sites, such as the hemolymph, higher detection of the pathogen might overwhelm the host’s immune response and concurrently result in terminal infection. In the empirical observation by Duneau et al. (2017), flies that succumbed to terminal infection exhibited a higher pathogen load upon death compared to the live flies. Further, Duneau et al. (2025) has confirmed with empirical observation and theoretical support that a host exhibiting a delayed immune response is susceptible to a high PLUD, which supports our model. The bifurcation of the infection trajectory possibly reflects a threshold until which the infection is stable, and once the host immunity is overwhelmed due to accumulated damage or the rapid outflow of bacteria, it results in the occurrence of terminal infection.

Studies examining infection outcomes in insects have primarily focused on bacterial load in the whole body (Duneau et al., 2017; Acuña-Hidalgo et al., 2022). However, our finer-scale spatial approach provides insight into the possible cause of these infection outcomes: The difference in the infection dynamics across body parts might be associated with a combination of events like circulatory and immune-specific (Sigle & Hillyer, 2016; Subasi et al., 2024). Future studies are needed to understand the spatial dynamics of infection outcomes based on immune differences in the body parts, and the migration and localized expression of AMPs with respect to the localization of the pathogen.

## Acknowledgements

We thank Jens Rolff, Brian P. Lazzaro, & Moria Chambers for thoughtful discussions. We thank Brian P. Lazzaro for the bacterial culture. We also thank Alisa Burkaltsava, Lina Ismail, and Julianna Yeliz Camurdas for help in the lab. Further, we would like to thank the Deutsche Forschungsgesellschaft for funding S. Armitage through Heisenberg Fellowship (AR 872/4-1 and AR 872/7-1). M. Menon was supported by the DFG through project AR 872/3-2, awarded to S. Armitage, and M. Franz was supported by the DFG (FR 3061/6-1) as part of the Research Unit FOR-2 ‘InsectInfect’.

---

**Figure S1.**
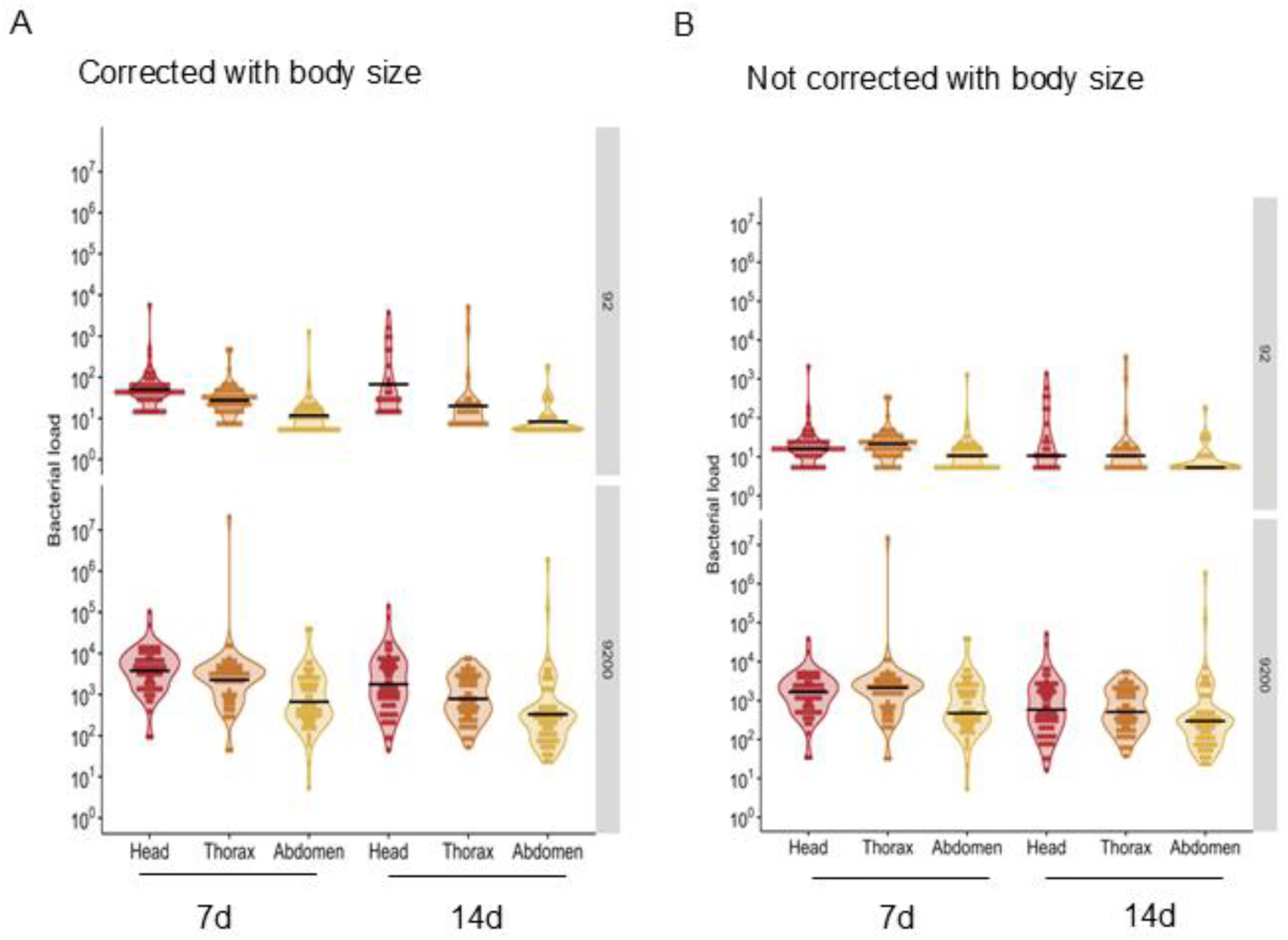
Bacterial load across all the body parts for ∼92 CFU and ∼9200 CFU doses during persistent infection. (complementary to Figure 3). The bacterial load is represented for two later time-points, 7d and 14d. A) Bacterial load, which is corrected for body part size. B) The bacterial load is not corrected for body part sizes. The thick black line indicates the mean. Each filled circle represents data points from an individual bacterial load.

**Figure S2.**
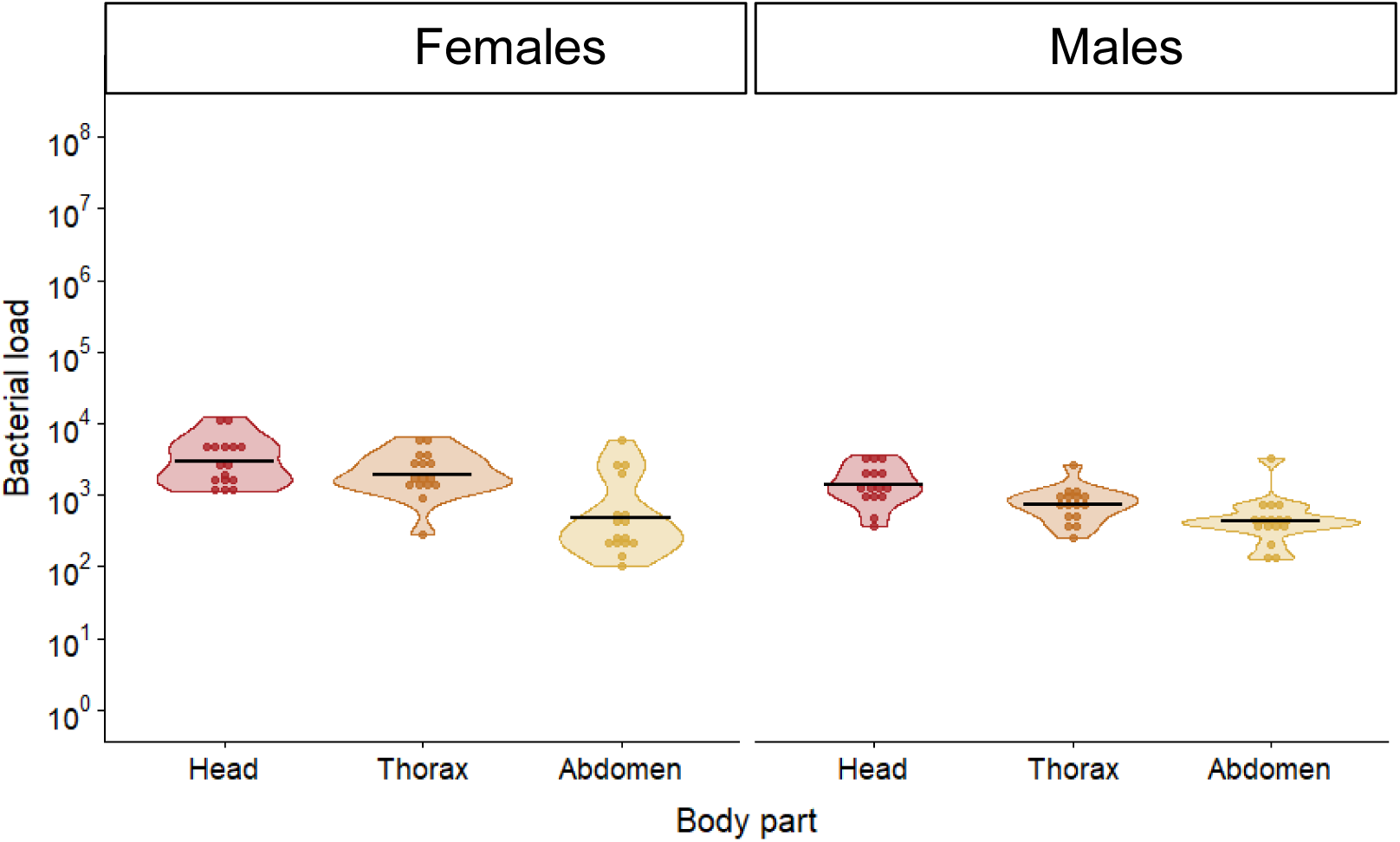
**Bacterial load in females and males during persistent infection**. The bacterial load was corrected for body size in males and females. This plot includes only day 7 post-injection in the thorax. The x-axis represents the body parts, and y-axis represents the bacterial load. The thick black line represents the mean. Each dot represents individual data points from a single fly’s body part. The experiment included a dose of ∼9200 CFU.

**Figure S3.**
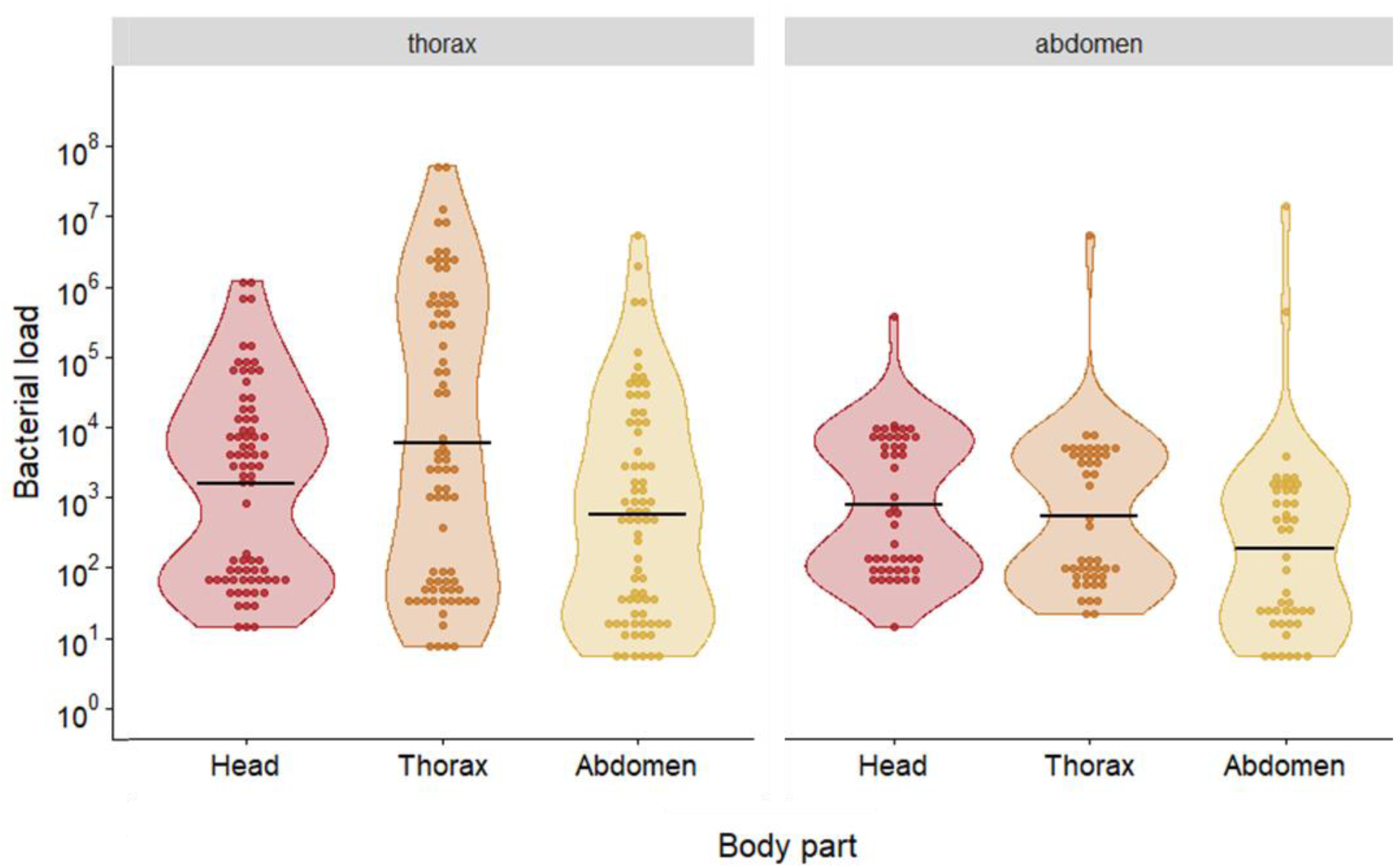
**Bacterial load during terminal infection corrected for body size**. This plot includes only day 1 post-infection. The x-axis represents the body parts, and the y-axis represents the bacterial load for both sites of injection, the abdomen and thorax. The thick black line represents the mean. Each dot represents individual data points from a single fly’s body part. The experiment included doses ∼92 CFU and ∼9200 CFU. Abdomen injection n = 137 individuals and thorax injection n = 225 individuals.

**Figure S4.**
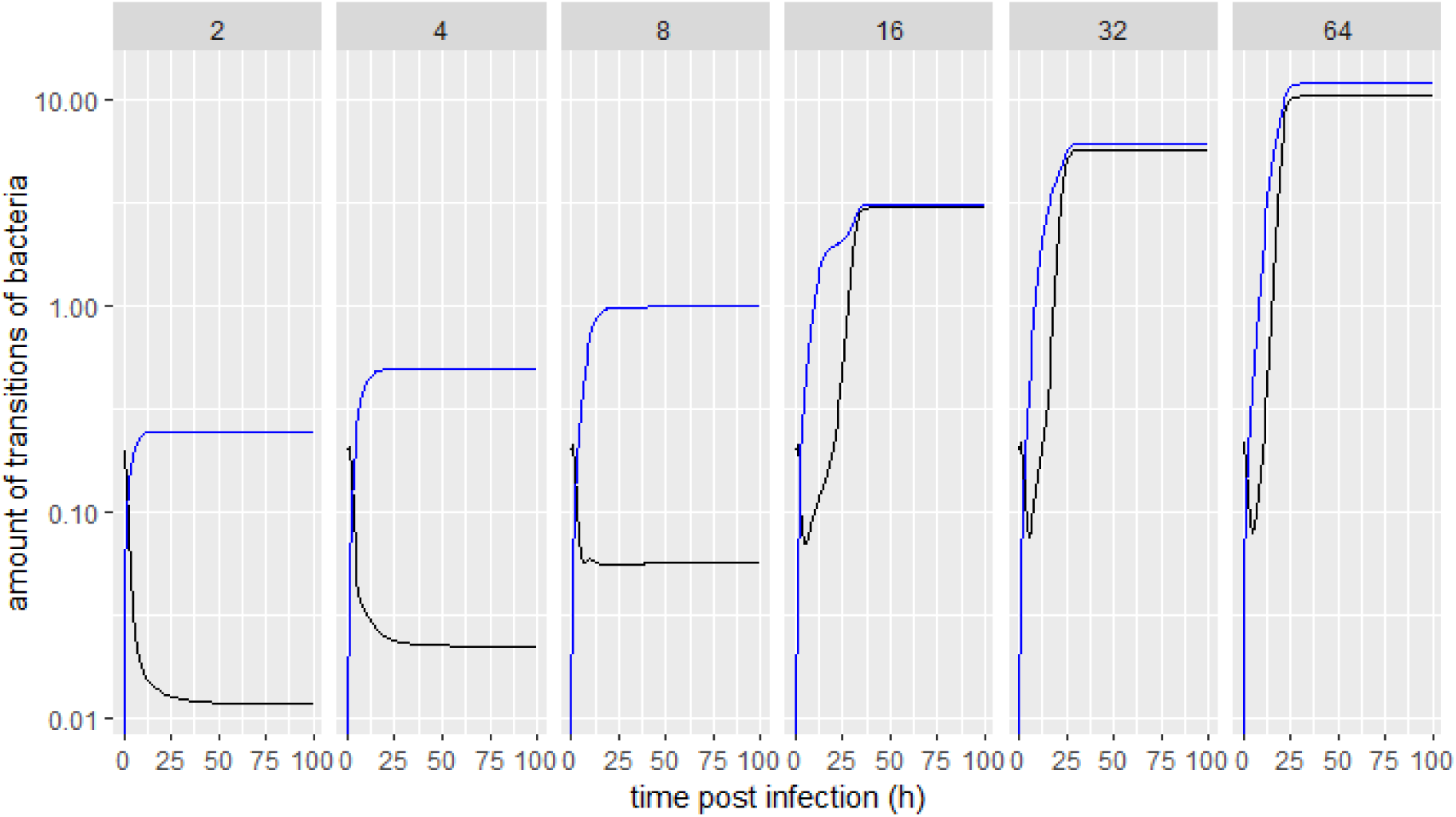
**Simulation results showing the transition of bacteria between tissues**. Simulated model dynamics depicting temporal changes in the number of bacteria that transition from the haemocoel to the protected tissue (black) and from the protected tissue to the haemocoel (blue), separately for different carrying capacities of the protected tissue *L*.

**Figure S5.**
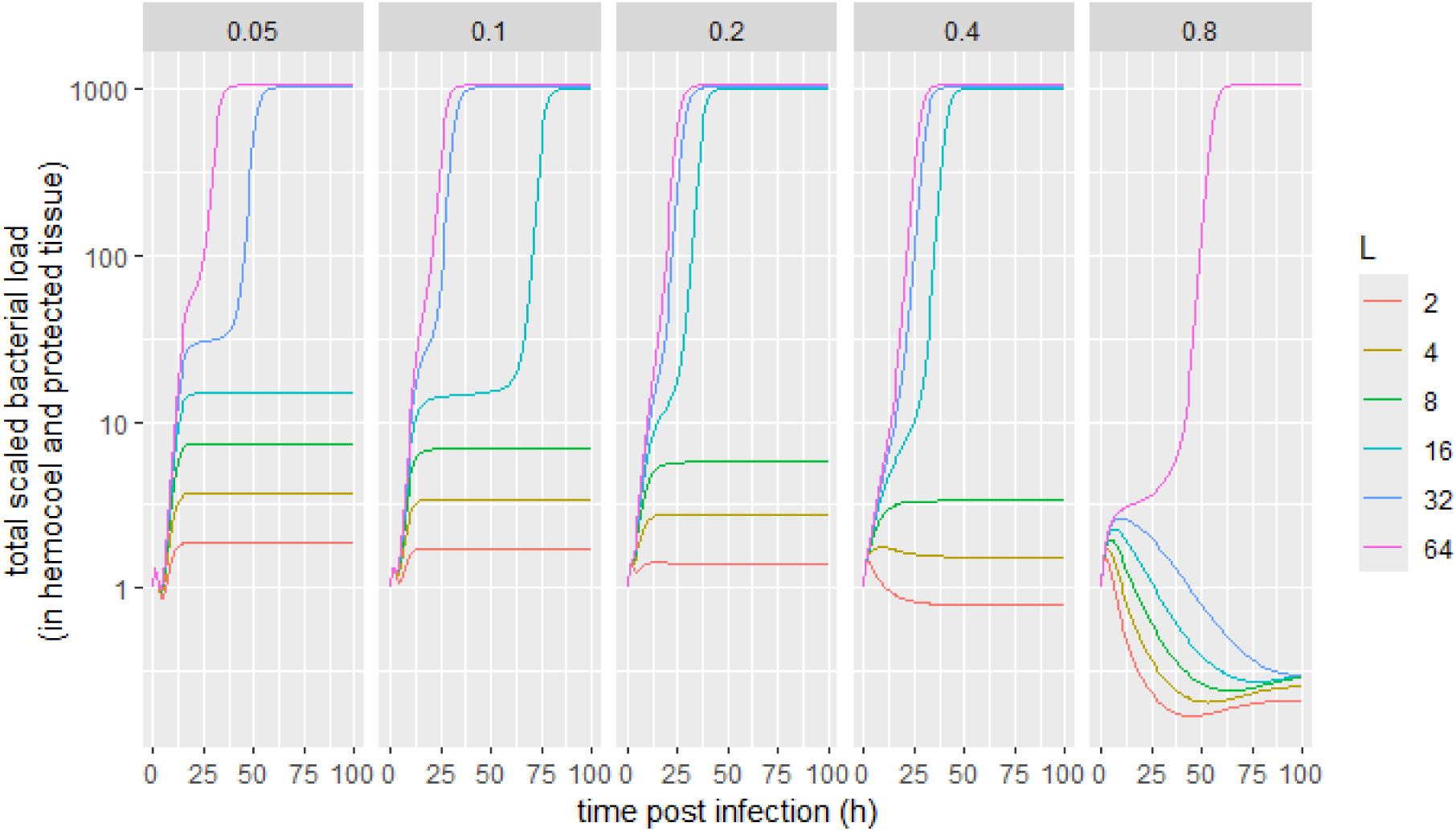
**Additional simulation results on the influence of transition rates on the emergence of bifurcating infections**. Simulated model dynamics depicting temporal changes in total scaled bacterial population size for different carrying capacities of the protected tissue *L*, separately for different transition rates *s*.

**Table S1.** Results of a linear mixed model with uncorrected bacterial load on day 7 and 14 as response variable (model 2). Bacterial load was not corrected for body size differences. The body part, site of injection, dose, and time post-infection were included as fixed effects, and replicate was included as a random effect. The body parts include head, thorax, and abdomen. Flies were injected through the thorax or abdomen with either ∼9200 CFU or ∼92CFU and were assayed at 7d and 14d post-infection. Statistically significant differences are given in bold.

| **Factor** | **Chisq** | **df** | ***p*** |
| --- | --- | --- | --- |
| Body part | 81.76 | 2 | **< 0.001** |
| Site of injection | 2.095 | 1 | **0.1478** |
| Dose | 1225.851 | 1 | **< 0.001** |
| Timepoint post-injection | 39.197 | 1 | **< 0.001** |
| Replicate | 72.57 | 4 | **< 0.001** |

**Table S2.** Tukey’s multiple comparison test for uncorrected data The post hoc tests for the difference in bacterial load among body parts for uncorrected data. Statistically significant differences are given in bold.

| **Contrast** | **T- test** | **df** | **p** |
| --- | --- | --- | --- |
| Abdomen- head | -7.760 | 581 | < 0.001 |
| Abdomen- thorax | -8.702 | 581 | < 0.001 |
| head- thorax | -0.321 | 581 | 0.94 |

**Table S3:** Post hoc comparison between the treatments for wounding and site of **inoculation for body size corrected data in terminal infection**. The table was generated using Fisher’s test to test the differences in high loads between the groups.

| **Treatment1** | **Treatment2** | ***p*** |
| --- | --- | --- |
| abd inj - thorax prick | abdomen injection | 0.494 |
| abd inj - thorax prick | thorax inj - abd prick | 0.090 |
| abd inj - thorax prick | thorax injection | 0.090 |
| abdomen injection | thorax inj - abd prick | 0.005 |
| abdomen injection | thorax injection | 0.005 |
| thorax inj - abd prick | thorax injection | 1.000 |

